# Optimising passive eDNA sampling: A theoretical framework for time-dependent eDNA accumulation

**DOI:** 10.64898/2026.08.17.745366

**Authors:** Hitoshi Araki, Masayuki K. Sakata

## Abstract

1. Environmental DNA (eDNA) methods are developing rapidly for ecological surveys, and passive eDNA sampling has emerged as a promising approach for integrating DNA signals over deployment time. However, how deployment duration affects the amount of detectable DNA retained by a sampler remains poorly understood.
2. Here, an analytical model was developed to examine how DNA input, degradation, finite substrate capacity and residual retention of degraded DNA shape passive eDNA accumulation. The model distinguishes detectable adsorbed DNA from degraded, non-detectable DNA that may remain on the substrate and continue to occupy capacity. The residual-retention parameter, *θ*, represents the fraction of degraded DNA that remains capacity-occupying, with *θ* = 0 corresponding to complete replacement and *θ* = 1 to complete non-replacement.
3. The model predicts three key behaviours. First, when degraded DNA does not occupy substrate capacity (*θ* = 0), detectable eDNA accumulates monotonically towards equilibrium, but equilibrium recovery increases less than proportionally with DNA input. Thus, passive-sampler measurements can compress quantitative differences in environmental DNA supply. Second, when degraded DNA remains capacity-occupying (*θ* > 0), detectable eDNA can reach a finite peak and subsequently decline. Higher DNA input increases peak yield but shifts the peak earlier, whereas greater substrate capacity increases peak yield and delays the peak. Third, under prolonged deployment with *θ* > 0, a higher-input condition can yield less detectable eDNA than a lower-input condition, reversing the expected input-rate ranking.
4. These results show that passive eDNA recovery can follow saturating, unimodal or intermediate dynamics depending on substrate capacity and post-adsorption DNA fate. Thus, retrieval time cannot be optimised by adjusting deployment duration alone. Although investigators can choose deployment duration and sampler design, including substrate capacity, optimisation also requires calibration or explicit assumptions about ambient DNA supply, DNA degradation rate and residual retention of degraded DNA.

## 1. Introduction

Environmental DNA (eDNA) analysis has become a widely used approach for detecting and monitoring aquatic organisms without the need for direct capture or visual observation. Since early demonstrations that aquatic macroorganisms can be detected using DNA recovered from water samples, eDNA has been increasingly applied to biodiversity assessment, rare species detection, invasive species surveillance, and conservation monitoring (Ficetola et al., 2008; Ruppert et al., 2019; Kanbe et al., 2023). Most aquatic eDNA studies have relied on active water sampling, in which a known volume of water is collected or filtered at a specific time and place. This approach provides a snapshot of the eDNA present in the sampled water volume (Yamamoto et al., 2016), but it can be sensitive to short-term temporal variation, spatial heterogeneity, and stochastic variation in DNA distribution (Beentjes et al., 2019; Troth et al., 2021; Hervé et al., 2022).

Passive eDNA sampling has recently attracted attention as an alternative or complementary approach to conventional water sampling (Chen et al., 2024; Sandré et al., 2026). In passive sampling, a substrate, membrane, or other collector is deployed in the environment for a defined period, during which environmental DNA is expected to adsorb onto or become retained within the sampler material. Recent studies have shown that eDNA can be passively collected using submerged membranes and other substrates, including positively charged nylon membranes and 3D-printed hydroxyapatite samplers, and that such approaches can recover biodiversity signals comparable to active filtration under some conditions (Bessey et al., 2021; Verdier et al., 2022; Chen et al., 2024). A recent review also emphasized that passive eDNA sampling is rapidly emerging as a promising complement or substitute for active sampling, while noting that convergence of methodological approaches remains limited (Sandré et al., 2026). The time-integrative property of passive samplers may be particularly useful when target DNA concentrations are low, when organisms are intermittently present, or when frequent active sampling is logistically difficult.

Despite these advantages, the interpretation of passive eDNA sampler-derived signals remains less straightforward than that of conventional water samples. The amount of DNA recovered from a passive sampler can be affected not only by ambient eDNA concentration or the rate at which DNA is supplied to the sampler, but also by processes occurring at the sampler surface, including DNA adsorption, retention, loss of molecular detectability, release, extraction efficiency, and possible saturation of substrate capacity. Previous studies have shown that eDNA degradation and persistence in aquatic environments are affected by environmental conditions, including temperature, microbial activity, and other physicochemical factors (Barnes et al., 2014; Collins et al., 2018). Moreover, eDNA does not occur as a single homogeneous entity; rather, it exists in multiple states, including dissolved, particle-associated, intracellular, and organellar forms, each of which may differ in its subsequent persistence and assay detectability (Mauvisseau et al., 2022). Passive sampler yield may therefore not scale linearly with environmental DNA supply or retrieval time. Indeed, recent field experiments have shown that the effects of submergence time vary among sampler materials and target taxa, with some combinations yielding less eDNA after longer than after intermediate deployments (von Ammon et al. 2025). However, the processes responsible for such non-monotonic recovery remain unclear.

A key unresolved issue is the relationship between molecular detectability and substrate occupancy after DNA has been retained by a sampler. One possibility is that degraded DNA is released from the substrate, thereby freeing capacity for newly supplied DNA. Under this assumption, detectable eDNA accumulation should approach a dynamic equilibrium determined by DNA input, DNA degradation, and substrate capacity. Alternatively, DNA may lose molecular detectability while remaining physically associated with the substrate surface or matrix. Because eDNA detection typically relies on amplification of target fragments by qPCR, ddPCR, or metabarcoding, DNA may become non-detectable if it is fragmented, chemically modified, or otherwise altered beyond the requirements of the target assay. If such non-detectable DNA continues to occupy finite substrate capacity, the amount of detectable DNA retained by the sampler may decline, even while total substrate occupation continues to increase. These alternative scenarios have fundamentally different implications for passive sampler deployment duration and quantitative interpretation.

The lower recovery observed after longer submergence times for some sampler– taxon combinations is consistent with several possible mechanisms, including continued occupation of the substrate by non-detectable DNA, desorption of previously retained DNA, and temporal variation in ambient DNA supply (von Ammon et al., 2025). Current empirical observations do not necessarily distinguish among these mechanisms. Nevertheless, explicitly separating molecular detectability from substrate occupancy provides a testable theoretical explanation for how deployment duration may affect eDNA recovery. Long deployments may be advantageous in low-DNA environments if accumulated DNA remains detectable or if capacity vacated by non-detectable DNA becomes available to newly supplied DNA. Conversely, if non-detectable DNA continues to occupy finite substrate capacity, detectable eDNA may decline during extended deployment. Substrate capacity may therefore determine not only the maximum amount of DNA retained but also the time window over which a sampler remains effective. A theoretical framework incorporating DNA input, degradation, and substrate capacity can thus clarify when passive eDNA sampling behaves as a cumulative sampling method and when it instead has a finite, yield-maximizing retrieval time.

In this study, a theoretical model for optimising passive eDNA sampling was developed and analytically solved to predict time-dependent eDNA accumulation on passive sampler substrates. The model introduces *θ* as the fraction of degraded DNA that remains on the substrate and continues to occupy substrate capacity. This formulation covers two limiting cases of post-adsorption DNA fate: complete replacement, in which DNA that becomes non-detectable no longer occupies substrate capacity (*θ* = 0), and complete non-replacement, in which all non-detectable DNA continues to occupy capacity (*θ* = 1). The model was used to examine how DNA input rate, degradation rate, residual retention of degraded DNA, and finite substrate capacity affect detectable eDNA yield over deployment time. By comparing these limiting cases and intermediate values of *θ*, this study identifies conditions under which passive eDNA sampling produces monotonic accumulation towards equilibrium or a unimodal response with a finite yield-maximizing retrieval time. It also examines whether finite capacity and prolonged deployment can distort proportional differences among DNA input conditions or reverse their expected rank order. The framework aims to support the design of passive sampler deployments and the quantitative interpretation of time-dependent passive eDNA data.

## 2. Materials and Methods

### 2.1 Conceptual framework

The framework was designed to describe how the amount of detectable DNA recovered from a passive sampler changes as a function of deployment time, environmental DNA supply, DNA degradation, residual retention of degraded DNA, and the finite DNA-holding capacity of the sampler substrate. A substrate-occupancy model was developed in which detectable adsorbed DNA can become degraded or non-detectable, and a fraction of this degraded DNA may retain on the substrate and continue to occupy capacity.

The model distinguishes two substrate-bound DNA states: detectable adsorbed DNA, *L*(*t*), and degraded, non-detectable DNA that remains capacity-occupying, *M*(*t*). The DNA input rate, *S*, represents the rate at which environmental DNA is supplied to and adsorbed by a fully vacant substrate. The degradation rate, *λ*, represents the rate at which detectable adsorbed DNA becomes degraded or non-detectable. The residual-retention parameter, *θ*, represents the fraction of degraded DNA that remains on the substrate and continues to occupy substrate capacity, where 0 ≤ *θ* ≤ 1. The substrate capacity, *K*, represents the maximum amount of DNA or DNA-derived material that can be accommodated by the sampler substrate.

This formulation includes two limiting cases of post-adsorption DNA fate. When *θ* = 0, degraded DNA does not remain capacity-occupying, and degradation frees substrate capacity for subsequent DNA capture. This case corresponds to complete replacement. When *θ* = 1, all degraded DNA remains on the substrate and continues to occupy capacity, corresponding to complete non-replacement. Intermediate values of 0 < *θ* < 1 represent partial residual retention of degraded DNA.

### 2.2 Substrate-occupancy model

Let *L*(*t*)denote the amount of detectable adsorbed DNA accumulated on the sampler at retrieval time *t*, measured as the elapsed time since sampler deployment. Let *M*(*t*)denote the amount of degraded, non-detectable DNA that remains on the substrate and continues to occupy substrate capacity. The temporal dynamics were described as

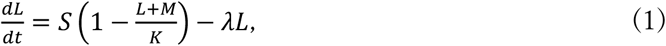

and

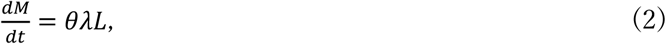

where *S* is the DNA input rate, *K* is the substrate DNA-holding capacity, *λ* is the degradation rate, and *θ* is the residual-retention fraction of degraded DNA. The term *S*(1 − (*L* + *M*)/*K*) represents DNA adsorption to vacant substrate capacity, whereas *λL* represents degradation or loss of molecular detectability of substrate-bound detectable DNA. Of the degraded DNA produced at rate *λL*, the fraction *θ* remains on the substrate and continues to occupy capacity. The initial conditions were *L*(0) = 0 and *M*(0) = 0, corresponding to a clean sampler at the start of deployment.

For constant *S*, *λ*, *θ*, and *K*, the model has an analytical solution. Setting *c* = *S*/*K* and

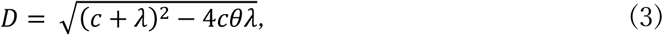

the two decay constants are

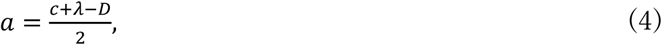

and

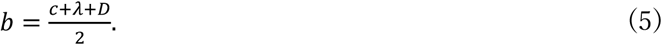

For *D* > 0, detectable DNA is given by

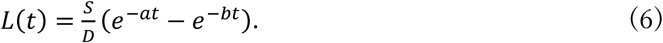

This expression includes complete replacement, complete non-replacement, and intermediate partial-retention cases within a single analytical framework.

### 2.3 Limiting cases

When *θ* = 0, degraded DNA does not remain capacity-occupying. In this complete-replacement case, Eq. (6) reduces to

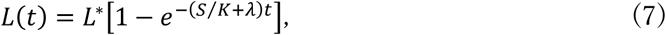

where

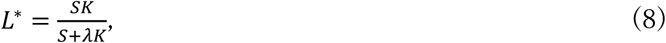

is the equilibrium amount of detectable DNA accumulated on the sampler. Thus, detectable DNA increases monotonically towards equilibrium.

When *θ* = 1, all degraded DNA remains on the substrate and continues to occupy capacity. In this complete non-replacement case, Eq. (6) reduces to

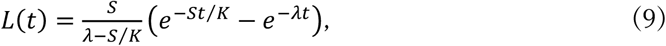

When *λ* ≠ *S*/*K*. If *λ* = *S*/*K*, the solution becomes

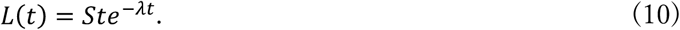

In contrast to the complete-replacement case, the complete non-replacement case predicts a unimodal trajectory in which detectable DNA increases to a maximum and then declines.

### 2.4 Peak retrieval time and maximum detectable DNA

For 0 < *θ* ≤ 1, detectable DNA reaches a finite maximum. When *D* > 0, the retrieval time that maximises detectable DNA is

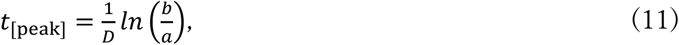

where a, b, and D are defined in Eqs. (3)-(5). The corresponding maximum amount of detectable DNA is

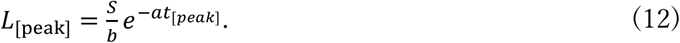

When *θ* = 0, no finite peak occurs; detectable DNA instead increases monotonically towards the equilibrium value *L*\*, as described above. In the special case of complete non-replacement with *θ* = 1 and *λ* = *S*/*K*, the peak occurs at

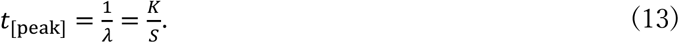

### 2.5 Analytical evaluation and parameter comparisons

All theoretical curves for both the basic and non-replacement models were generated by direct evaluation of their analytical solutions. The basic accumulation and non-replacement models were evaluated over the specified retrieval-time range using the baseline parameter values of *S* = 1.0, *λ* = 0.03, and *K* = 40. The residual-retention fraction θ was varied between 0 and 1 to examine the transition from complete replacement (*θ* = 0) to complete non-replacement (*θ* = 1), with *θ* = 0.5 representing partial residual retention of degraded DNA. The effects of DNA input rate, degradation rate, retention of degraded DNA, and substrate capacity were examined by varying *S*, *λ,* θ, and *K*, respectively, while holding the other parameters constant.

For comparisons between two DNA input rates, *S_i_* and *S_j_*, the difference in detectable DNA recovery in the non-replacement model was calculated as

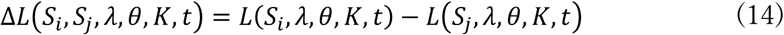

and the recovery ratio was calculated as

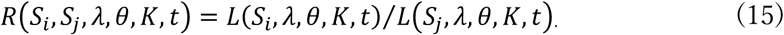

When *S_i_* > *S_j_*, positive values of Δ*L*, or values of *R* > 1, indicate that treatment *S_i_* produced more detectable DNA than treatment *S_j_*. as expected from their input rates. Negative values of Δ*L*, or values of *R* < 1, indicate a reversal of the original input-rate ranking. Parameter-space plots and time-series comparisons were produced by evaluating Eqs. (6), (8), and (14), and (15) directly over the specified combinations of *S*, *λ*, θ, *K*, and retrieval time.

Phase-diagram analyses were conducted to examine how the model predicts yield-maximising retrieval timing and maximum detectable DNA recovery across parameter space. To allow comparison among parameter combinations with different input rates and substrate capacities, peak timing and peak yield were expressed in dimensionless form. The dimensionless peak retrieval time, (*S*⁄*K*)*t*_[peak]_, and the peak detectable DNA yield relative to substrate capacity, *L*_[peak]_/*K*, were calculated by direct evaluation of the analytical solutions over a range of residual-retention fractions, θ, and dimensionless degradation parameters, *λK*/*S*. The dimensionless degradation parameter represents the ratio of the substrate-filling timescale (*K*/*S*) to the DNA degradation timescale (1/*λ*). All analytical evaluation and parameter comparisons were conducted using R ver. 4.6.0. Graphical outputs were generated using the R package “ggplot2”.

## 3. Results

### 3.1 Saturating and unimodal accumulation dynamics

The analytical solution showed that the residual-retention fraction of degraded DNA, *θ*, qualitatively altered the temporal dynamics of detectable eDNA accumulation (Fig. 1). Under the complete replacement case (*θ* = 0), detectable eDNA increased monotonically towards an equilibrium because degradation released substrate capacity for subsequent DNA capture. With the baseline parameter values *S* = 1.0, *λ* = 0.03, and *K* = 40, the equilibrium amount of detectable eDNA, *L*\*, was 18.18, as calculated from Eqn 8.

**Fig. 1.**
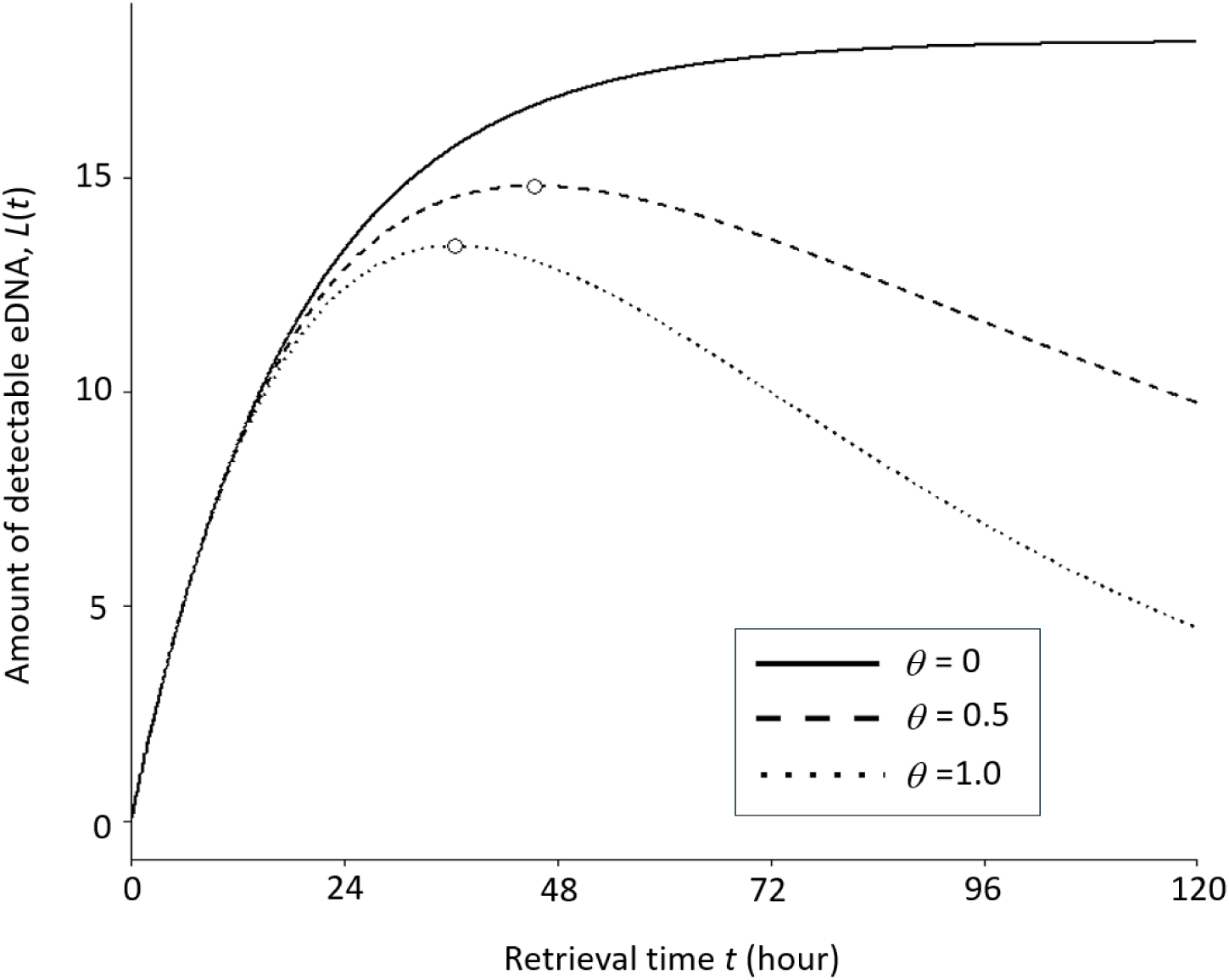
Effects of parameter changes on eDNA accumulation predicted by the model under an assumption of complete replacement (*θ* = 0). Detectable eDNA, *L*(*t*), was calculated with baseline values *S* = 1.0, *λ* = 0.03, and *K* = 40. One parameter was doubled at a time while the others were fixed. The thin solid line shows the baseline condition; thick solid, dotted, and dashed lines show increased *S*, *λ*, and *K*, respectively.

In contrast, when degraded DNA was partially or fully retained on the substrate (*θ* > 0), detectable eDNA followed unimodal trajectories with finite maximums. Under otherwise identical parameter values, partial residual retention (*θ* = 0.5) produced a peak of *L*_[peak]_ = 14.8 at *t*_[peak]_ = 45.4h, whereas complete non-replacement (*θ* = 1.0) produced a lower and earlier peak of *L*_[peak]_ = 13.4 at *t*_[peak]_ = 36.5h (Fig. 1). Thus, increasing residual retention of degraded DNA shifted detectable eDNA accumulation from monotonic saturation towards earlier peak timing and stronger post-peak decline.

### 3.2 Substrate saturation compresses input-rate differences

Even under the complete replacement case (*θ* = 0), detectable eDNA recovery was not proportional to DNA input rate because of finite substrate capacity (Fig. 2). Doubling the DNA input rate from *S* = 1.0 to 2.0 increased *L*\* from 18.18 to 25.00, representing a 1.38-fold rather than twofold increase. A four-fold increase of input rate, from *S* = 1.0 to 4.0, increased *L*\* only to 30.77, corresponding to a 1.69-fold increase. This indicates that finite substrate capacity compressed quantitative differences in DNA input even when detectable eDNA accumulated monotonically.

**Fig. 2.**
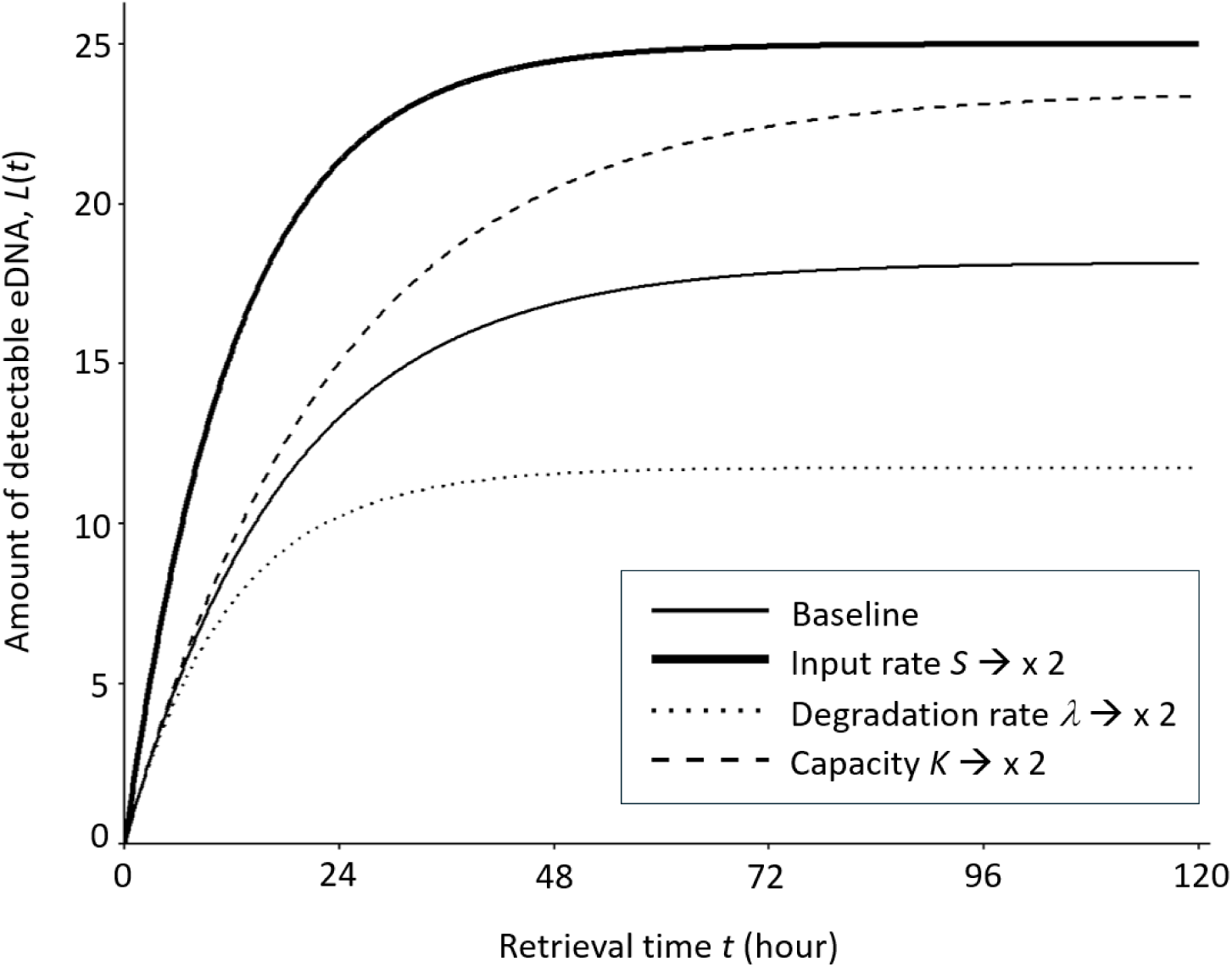
Comparison of analytical predictions from the model under identical parameter values of *S* = 1.0, *λ* = 0.03, and *K* = 40. Solid, dashed, and dotted lines indicate detectable eDNA, *L*(*t*), for *θ* = 0, 0.5, and 1.0, respectively. Open circles indicate the predicted peaks for *θ* = 0.5 (*L*_[peak]_ =14.80, *t*_[peak]_ =45.44h) and *θ* = 1.0 (*L*_[peak]_ =13.40, *t*_[peak]_ =36.46h).

Doubling the degradation rate from *λ* = 0.03 to 0.06 reduced the equilibrium amount to *L*^∗^ = 11.76, whereas doubling substrate capacity from *K* = 40 to 80 increased it to *L*^∗^ = 23.53 and slowed the approach to equilibrium. Together, these results indicate that detectable eDNA recovery under complete replacement is governed by the combined effects of DNA input rate, degradation rate and finite substrate capacity, rather than by DNA input rate alone.

### 3.3 Parameter effects on peak yield and retrieval time under non-replacement

Under the complete non-replacement case (*θ* = 1.0), detectable eDNA showed a unimodal trajectory, and changes in DNA input rate, degradation rate and substrate capacity altered both peak yield and peak timing (Fig. 3). Increasing the DNA input rate increased the maximum amount of detectable eDNA but shifted the peak to earlier retrieval times (Fig. 3a). With *λ* = 0.03 and *K* = 40, increasing *S* from 1.0 to 2.0 and 4.0 increased *L*_[peak]_ from 13.4 to 18.6 and 23.9, respectively, while shifting *t*_[peak]_ from 36.5 h to 25.5 h and 17.2 h.

**Fig. 3.**
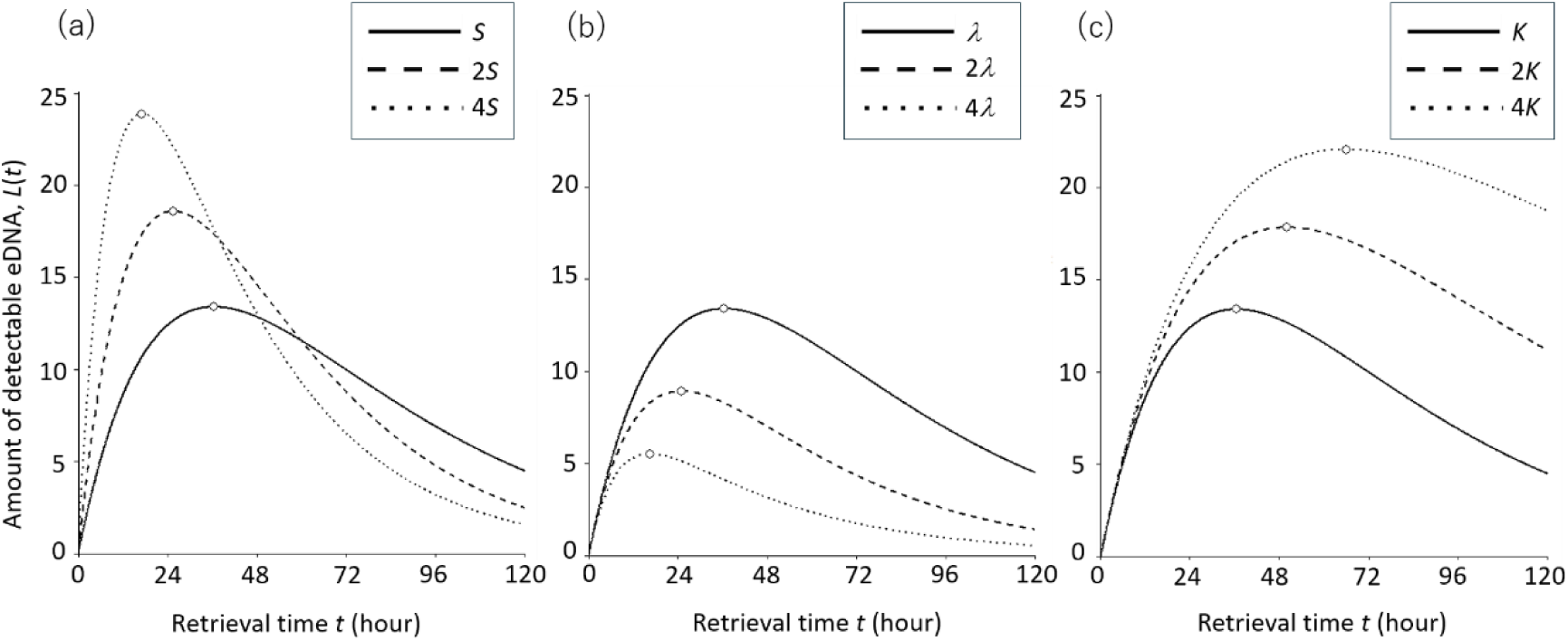
Effects of parameter variation on detectable eDNA accumulation in the model with *θ* = 1.0. Detectable eDNA, *L*(*t*), was calculated with baseline values *S* = 1.0, *λ* = 0.03, and *K* = 40, and one parameter was varied in each panel: (a) input rate *S*, (b) degradation rate *λ*, and (c) substrate capacity *K*. Solid, dashed, and dotted lines indicate baseline, twofold, and fourfold values, respectively. Open circle indicates the predicted peak for each case.

Increasing the degradation rate reduced the peak amount of detectable eDNA and also shifted the peak earlier (Fig. 3b). With *S* = 1.0and *K* = 40, increasing *λ* from 0.03 to 0.06 and 0.12 reduced *L*_[peak]_ from 13.4 to 8.9 and 5.5, respectively, while shifting *t*_[peak]_ from 36.5 h to 25.0 h and 16.5 h. By contrast, increasing substrate capacity increased the peak amount and delayed peak timing (Fig. 3c). With *S* = 1.0 and *λ* = 0.03, increasing *K* from 40 to 80 and 160 increased *L*_[peak]_ from 13.4 to 17.8 and 22.1, respectively, while shifting *t*_[peak]_ from 36.5 h to 50.0 h and 66.0 h. Together, these results show that the yield-maximising retrieval time under non-replacement is parameter dependent. Higher DNA input and higher degradation rates shorten the optimal retrieval window, whereas greater substrate capacity extends it.

Phase-diagram analyses further summarized the combined effects of the residual-retention fraction, θ, and the dimensionless degradation parameter, *λK*/*S*, on peak retrieval time and peak detectable DNA yield (Fig. 5). The dimensionless peak retrieval time decreased with increasing θ and *λK*/*S* (Fig. 5a), indicating that greater retention of degraded DNA and faster loss of detectability both advanced the timing of maximum detectable DNA recovery. In contrast, peak detectable DNA yield was influenced primarily by *λK*/*S*, decreasing progressively as*λK*/*S* increased, whereas the effect of θ was comparatively modest (Fig. 5b). Together, these phase diagrams show that yield-maximising retrieval timing is particularly sensitive to residual retention of degraded DNA, whereas peak yield is governed mainly by the balance between degradation and substrate-filling rate.

### 3.4 Time-dependent reversal of input-rate rankings

Under non-replacement dynamics, differences in detectable eDNA recovery between input treatments depended strongly on input rate, degradation rate, and retrieval time (Fig. 4). This behaviour is illustrated here using the complete non-replacement case (*θ* = 1.0). At the baseline degradation rate of *λ* = 0.03 and substrate capacity of *K* = 40, Δ*L* was initially positive for all comparisons in which *S_i_* > *S_j_* = 1.0, indicating that greater DNA input initially produced greater detectable accumulation. However, Δ*L* declined with retrieval time and eventually became negative, demonstrating a reversal of the original input-rate ranking. This transition from positive to negative Δ*L* occurred earlier as *S_i_* increased, because higher input rates produced earlier peaks and therefore earlier post-peak declines.

**Fig. 4.**
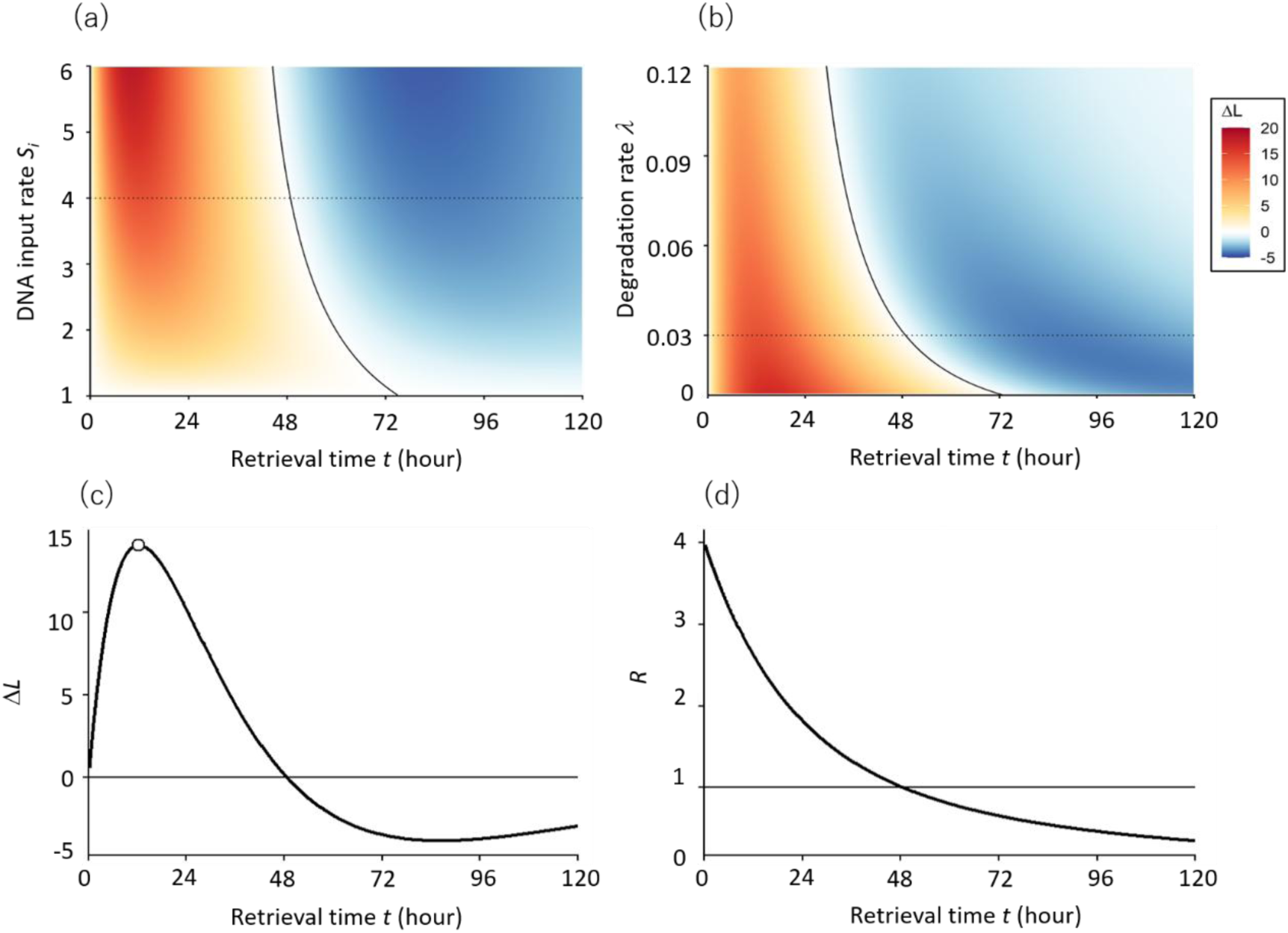
Retrieval-time effects on detectable eDNA accumulation under the complete non-replacement case (*θ* = 1.0). Calculations were performed with *K* = 40. (a) Δ*L* across input rate *S_i_* and retrieval time *t*, calculated using Eqn 14 with *S_j_* = 1.0 and *λ* = 0.03. (b) Δ*L* across degradation rate *λ* and retrieval time *t*, with *S_i_* = 4.0 and *S_j_* = 1.0. (c) Temporal change in Δ*L* for *S_i_* = 4.0, *S_j_* = 1.0, and *λ* = 0.03. (d) Temporal change in the recovery ratio, *R*, with the same parameters. Blue and red in panels (a) and (b) indicate positive and negative Δ*L*, respectively; black contours indicate Δ*L* = 0. Dotted horizontal lines in panels (a) and (b) indicate the parameter values used in panels (c) and (d). Under the conditions shown in panels (c) and (d), Δ*L*, the difference in detectable eDNA between the two input rates (*S_i_* = 4.0 and *S_j_* = 1.0), reaches its maximum at *t* = 12.4 (indicated by an open circle), and crosses zero at *t* = 48.8.

The timing of rank reversal also depended on the degradation rate (Fig. 4b). For the comparison between the fourfold input treatment (*S_i_* = 4.0) and the baseline treatment (*S_j_* = 1.0), Δ*L* remained positive for a longer period when *λ* was low. As *λ* increased, the zero contour shifted toward shorter retrieval times, indicating that faster loss of molecular detectability advanced the reversal of the input-rate ranking.

The temporal trajectory under the baseline conditions of *λ* = 0.03 and *K* = 40 illustrated this reversal directly (Fig. 4c, d). The fourfold input treatment initially produced substantially more detectable DNA than the baseline treatment, and Δ*L* reached its maximum at *t* = 12.4h. Thereafter, Δ*L* declined and crossed zero at *t* = 48.80h. After this crossover time, the treatment with fourfold greater DNA input yielded less detectable DNA than the baseline treatment.

The recovery ratio showed the same transition in relative recovery (Fig. 4d). Immediately after deployment, *R* approached the input-rate ratio of 4, reflecting the initial proportional effect of DNA input. The ratio subsequently declined, crossed *R* = 1 at *t* = 48.80, and remained below 1 thereafter. At *t* =120h, *R* was 0.35, indicating that detectable DNA under the fourfold input treatment was only approximately 35% of that under the baseline treatment. Thus, when degraded DNA remains capacity-occupying, prolonged retrieval times can not only compress differences in DNA input but also reverse their apparent rank order. This reversal was most clearly expressed under the complete non-replacement case (*θ* = 1.0).

## 4. Discussion

The analytical solutions developed in this study show that passive eDNA recovery can follow either saturating or unimodal dynamics, depending on the residual retention of degraded DNA on the sampler substrate (Fig. 1). When degraded DNA no longer occupies substrate capacity (*θ* = 0), detectable eDNA accumulates monotonically towards equilibrium. However, even in this complete replacement case, equilibrium recovery is not proportional to DNA input, indicating that passive-sampler measurements can compress quantitative differences among environments with different DNA supplies. When degraded DNA remains capacity-occupying (*θ* > 0), detectable eDNA reaches an analytically defined maximum and subsequently declines. In the complete non-replacement case (*θ* = 1), this decline occurs even as total substrate occupancy continues to increase towards capacity. Prolonged deployment can therefore reverse the rank order of detectable eDNA among input treatments, such that a site with greater DNA supply yields less detectable eDNA than a site with lower DNA supply. These results indicate that retrieval time is not merely a determinant of total eDNA yield, but also a determinant of the quantitative relationship between passive-sampler recovery and environmental DNA input.

### 4.1 Passive eDNA accumulation is nonlinear and parameter dependent

When degraded DNA no longer occupied substrate capacity (*θ* = 0), degradation effectively released capacity for subsequent DNA capture. Detectable eDNA therefore increased rapidly during early deployment and gradually approached an equilibrium determined by DNA input rate, degradation rate and substrate capacity (Fig. 2). This pattern resembles the integrative-to-equilibrium uptake kinetics described for conventional aquatic passive samplers, which commonly exhibit an initial integrative phase followed by slower uptake as exchange approaches equilibrium (Vrana et al., 2005). It also provides a potential mechanistic interpretation of time-dependent eDNA capture on submerged membranes and DNA-binding substrates, for which rapid capture and material-dependent temporal responses have been reported (Bessey et al., 2021; Verdier et al., 2022).

Even under this monotonic complete replacement case, however, sampler recovery was not proportional to eDNA input. The equilibrium amount, *L*^∗^ = *SK*/(*S* + *λK*), is a saturating rather than linear function of input rate. Under the parameter values used in Fig. 2, increasing *S*from 1.0 to 2.0 and 4.0 increased *L*^∗^from 18.18 to 25.00 and 30.77, respectively, corresponding to only 1.38-fold and 1.69-fold increases despite twofold and fourfold increases in input rate. Finite capacity therefore compressed differences among input rates as substrate occupancy increased. The complete replacement case preserved the rank ordering of DNA inputs, but underestimated their relative magnitude, indicating that substrate saturation can affect quantitative interpretation even when detectable eDNA does not decline through time.

A different trajectory emerged when degraded DNA remained, at least partly, on the substrate and continued occupying capacity (*θ* > 0). In these non-replacement cases, detectable eDNA initially accumulated but subsequently declined as capacity became increasingly occupied by degraded, non-detectable DNA or DNA-derived material (Fig. 1). The complete non-replacement case (*θ* = 1) produced the strongest form of this behaviour: detectable eDNA reached a finite maximum and subsequently declined, whereas total substrate occupancy continued to approach *K*. Increasing DNA input rate or degradation rate shifted the peak to earlier retrieval times, whereas increasing substrate capacity delayed the peak and increased peak yield (Fig. 3). Thus, non-replacement dynamics create a transient sampling window in which both the amount of detectable eDNA and the yield-maximising retrieval time are parameter dependent.

Previous passive eDNA studies have shown that longer submergence does not necessarily produce continued increases in DNA recovery or taxonomic detection, consistent with rapid capture, saturation or diminishing returns (Bessey et al., 2021, 2022). More recently, von Ammon et al. (2025) reported sampler- and taxon-specific cases in which eDNA yield was lower after longer than after intermediate submergence times. These observations provide empirical support for the possibility of non-monotonic recovery during passive sampling. However, because detectable DNA, degraded substrate-associated DNA and total occupied capacity were not quantified separately, the observed declines cannot be attributed specifically to residual occupation by degraded DNA. They may also reflect degradation, desorption, temporal variation in ambient DNA supply or other field processes. The unimodal trajectory predicted when *θ* > 0should therefore be regarded as a mechanistically explicit hypothesis that is consistent with some empirical observations but remains to be directly tested.

These results are relevant to the growing use of submerged membranes, mineral substrates and other passive materials for aquatic eDNA collection (Bessey et al., 2021; Verdier et al., 2022). Such approaches can effectively recover biodiversity signals, but accumulated DNA quantities cannot be interpreted without considering adsorption, degradation, release, residual retention and deployment duration. The absence of continued gains, or an apparent decline during prolonged deployment, may result from several distinct processes, including equilibrium-like saturation, declining molecular detectability, desorption or changing environmental DNA supply. Distinguishing among these explanations will require time-series measurements that separately quantify detectable DNA and total substrate-associated DNA or DNA-derived material.

Phase-diagram analyses across the model revealed that the parameters governing retrieval timing and maximum detectable DNA recovery differed (Fig. 5). Increasing the residual-retention fraction (*θ*) substantially advanced the yield-maximising retrieval time, but had only a modest effect on peak detectable DNA yield. In contrast, the dimensionless degradation parameter (*λK/S*) strongly affected both the timing and magnitude of peak recovery, with particularly pronounced effects on peak yield. These results suggest that different biological and physicochemical processes control when a passive sampler should be retrieved and how much detectable DNA can be recovered at the peak. Consequently, optimisation of deployment duration requires information not only on substrate capacity, but also on DNA degradation rate and the extent to which degraded DNA continues to occupy substrate sites.

**Fig. 5.**
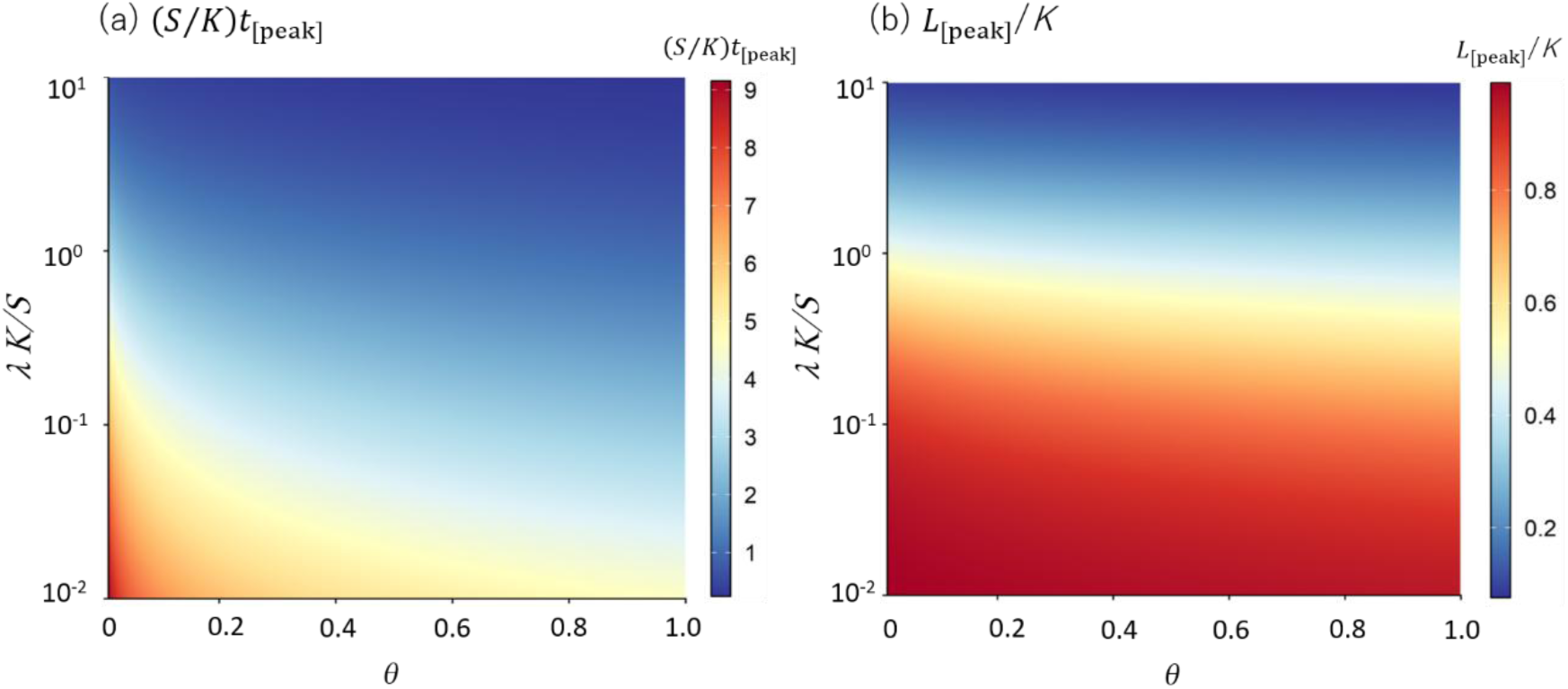
Parameter-space predictions of passive eDNA retrieval timing and peak recovery. Phase diagrams show the effects of the residual-retention fraction, θ, and the dimensionless degradation parameter, *λK*/*S*, on passive eDNA recovery. The parameter *λK/S* compares detectable DNA loss with substrate filling. (a) Dimensionless peak retrieval time, (*S*⁄*K*)*t*_[peak]_. (b) Peak detectable DNA yield relative to substrate capacity, *L*_[peak]_/*K*. Together, these diagrams show how degraded-DNA retention and the degradation–filling balance determine optimal retrieval timing and maximum detectable DNA recovery.

### 4.2 Irreversible occupation can obscure quantitative interpretation

The key mechanism underlying non-replacement dynamics is the decoupling of physical substrate occupation from analytical detectability. DNA may remain associated with the sampler after fragmentation or loss of the assay-specific target region has rendered it undetectable. A sampler can therefore become increasingly occupied by DNA-derived material while yielding decreasing quantities of amplifiable target DNA.

This decoupling creates ambiguity in endpoint measurements. A low qPCR, ddPCR or metabarcoding signal may indicate low environmental DNA supply, but it may also result from rapid degradation, limited effective capacity or retrieval after the detectable signal has passed its maximum. These possibilities cannot be distinguished from a single retrieval time. Such ambiguity is plausible because environmental DNA occurs in dissolved, particle-associated, intracellular and organellar forms that differ in adsorption, persistence and molecular detectability (Mauvisseau et al., 2022).

Under complete non-replacement (*θ* = 1), this decoupling is especially clear: total substrate occupancy depends on DNA input rate and substrate capacity, but not on the degradation rate. Degradation therefore changes how occupied capacity is partitioned between detectable and non-detectable DNA, rather than how rapidly the substrate becomes physically occupied. More generally, when degraded DNA remains capacity-occupying (*θ* > 0), higher DNA input can accelerate both detectable DNA accumulation and subsequent occupation by degraded, non-detectable DNA. Consequently, although higher input produces greater detectable recovery early in deployment, it can yield lower recovery than a lower-input condition after the high-input signal has peaked and declined (Fig. 4). Thus, complete replacement compresses input differences but preserves their ordering, whereas non-replacement dynamics can both compress and reverse them.

This prediction has implications for sampler design. Strong adsorption may improve short-term capture, but maximum binding strength may not optimise long-term molecular recovery if degraded molecules remain attached and continue to occupy capacity. Effective substrates may therefore require a balance among capture, retention and turnover, rather than simply the highest possible adsorption affinity.

### 4.3 Deployment duration and practical implications

The unimodal trajectories predicted under non-replacement dynamics imply a yield-maximising retrieval time, defined as the deployment duration at which detectable eDNA reaches its maximum. This retrieval time depended on DNA input rate, degradation rate, residual retention of degraded DNA and substrate capacity (Fig. 3). Increasing DNA input produced a faster initial rise and a higher peak, but shifted the peak earlier. Increasing the degradation rate reduced peak height and shortened the period during which detectable DNA remained elevated. By contrast, increasing substrate capacity increased peak recovery and delayed the peak by extending the interval before substrate occupation became limiting. However, greater capacity did not eliminate post-peak decline when degraded DNA remained capacity-occupying.

The analytical expressions for *t*_[peak]_ and *L*_[peak]_ show that the yield-maximising retrieval time is determined by the balance among substrate filling, molecular degradation and residual retention. Higher DNA input fills vacant substrate capacity more rapidly and advances the peak, whereas larger substrate capacity delays it. Faster degradation also advances the peak because detectable DNA is transferred more rapidly into a non-detectable state. When a fraction of this degraded DNA remains capacity-occupying (*θ* > 0), the yield-maximising retrieval time becomes a mechanistic property of DNA supply, substrate capacity and post-adsorption DNA fate, rather than an arbitrary feature identified from simulated curves.

Substrate capacity should be interpreted as an effective, material-specific property rather than a fixed universal constant. Passive sampler materials may differ in surface area, pore structure, mineral composition, charge properties, hydrophobicity and biofilm development, all of which can alter the number and accessibility of DNA-binding sites (Verdier et al., 2022; Sandré et al., 2026). In the present framework, these differences are represented by *K*, but they may also affect adsorption, retention and the residual-retention fraction *θ*. Materials with higher effective capacity may delay substrate limitation and extend the yield-maximising retrieval time, whereas low-capacity or rapidly saturated materials may peak earlier and become more susceptible to post-peak decline. Retrieval timing should therefore be calibrated for each sampler material.

This issue is particularly important for rare-species monitoring. Extended deployment is often expected to improve detection by integrating weak signals over time. However, passive samplers are exposed not only to target DNA but also to abundant background DNA from common organisms, microorganisms and extracellular material. If this background DNA occupies finite binding sites, it may progressively reduce the capacity available for rare target DNA. Under non-replacement dynamics, background DNA may continue to occupy sites even after becoming analytically undetectable. Prolonged deployment may therefore fail to improve rare-species detection and could reduce the opportunity for newly arriving target DNA to bind.

Practical calibration should use staggered retrieval times spanning the expected rising, peak and declining phases. Paired active-water samples could provide independent estimates of ambient DNA supply, while repeated passive-sampler retrievals would identify whether accumulation is monotonic, saturating or unimodal. Calibration across total DNA loads, degradation conditions and substrate types would help determine whether background DNA and material-specific capacity alter the optimal sampling window.

Deployment duration could then be selected according to the study objective, such as maximising detection probability, comparing DNA quantities among sites or integrating temporal exposure.

### 4.4 Implications for metabarcoding and abundance inference

Finite substrate capacity has different implications for among-sample quantitative comparisons and within-sample taxonomic composition. Under complete replacement (*θ* = 0), saturation caused absolute recovery to increase less than proportionally with DNA input. Comparisons among sites, seasons or deployment periods may therefore underestimate the true magnitude of differences in environmental DNA supply. Under non-replacement dynamics (*θ* > 0), this relationship also depended on retrieval time and could even reverse after high-input samples had passed their peaks.

Within a single sampler, taxonomic proportions could theoretically remain unchanged if DNA from all taxa entered simultaneously, competed for the same sites and shared identical adsorption and degradation properties. In practice, however, taxa may differ in DNA fragment-size distribution, physical state, adsorption affinity, degradation rate and accessibility to substrate surfaces. Subsequent laboratory and bioinformatic processes, including extraction, primer binding, PCR amplification, sequencing and sequence processing, can introduce additional taxon- and sequence-specific biases. Amplicon sequence variant proportions may therefore differ from both substrate-bound DNA ratios and environmental input ratios.

A multispecies extension of the present framework would be useful for examining how DNA sources compete for finite substrate capacity. Such a model could evaluate whether saturation merely compresses input ratios or whether taxon-specific adsorption, degradation and residual retention can distort, or even reverse, relative representation through time. Empirical evaluation would require mixed-DNA experiments with known taxon-specific inputs, combined with deployments across different total DNA concentrations and retrieval times. Passive-sampler metabarcoding may provide useful indices of relative DNA supply when the relevant taxa behave similarly, but read proportions should not automatically be interpreted as relative organism abundance.

The ecological meaning of the input parameter *S*also requires caution. In the present framework, *S*represents the effective rate at which DNA is supplied to and captured by the sampler, rather than numerical abundance or biomass directly. Quantitative eDNA signals may depend jointly on organism abundance, individual body size and population size structure. Yates et al. (2025) proposed an allometric framework linking eDNA to both numerical abundance and biomass, emphasising that neither quantity alone necessarily determines DNA production. For passive sampling, this organism-to-eDNA relationship is further modified by transport, adsorption, degradation, saturation and deployment duration. Accumulated DNA should therefore not be interpreted as proportional to abundance or biomass without system-specific calibration.

### 4.5 Experimental tests, model limitations, and future extensions

The contrasting predictions of the limiting cases are experimentally testable. A monotonic increase followed by a plateau would support complete replacement (*θ* = 0), in which DNA loss releases substrate capacity, whereas a detectable signal that peaks and declines while total substrate-associated DNA continues to increase would support complete or near-complete non-replacement dynamics. Experiments should therefore quantify both amplifiable target DNA and total substrate-bound DNA across retrieval time. qPCR or ddPCR could measure detectable target DNA, whereas fluorometric assays, DNA staining, fragment-size analysis or sequencing-based approaches could help characterise total and partially degraded DNA.

Sequential-exposure experiments would provide a direct test of substrate-site replacement. For example, a substrate could first be exposed to one identifiable DNA source, incubated under degradation-promoting conditions and then exposed to a second source.

Reduced capture of the second source would support persistent site occupation, whereas normal capture would indicate release, replacement or turnover. Experiments using multiple input concentrations would also test whether finite capacity compresses recovery ratios.

The framework is deliberately simplified. DNA input rate, degradation rate, residual retention and substrate capacity were treated as constant, target DNA was considered in isolation, and the substrate was represented as a homogeneous pool of identical binding sites. In natural systems, eDNA supply varies through time, adsorption and retention may depend on fragment size, surface chemistry, flow, water chemistry, biofilm development and competition with non-target DNA or particles, and degradation likely proceeds continuously rather than through discrete detectable and non-detectable states (Strickler et al., 2015; Shogren et al., 2017; Mauvisseau et al., 2022; Snyder et al., 2023; Jo, 2023). The complete replacement and complete non-replacement cases should therefore be regarded as mechanistic endpoints within a broader continuum of passive-sampler behaviour.

Future extensions could incorporate turnover of degraded DNA, heterogeneous binding sites, progressive degradation, time-varying inputs, multispecies competition and explicit observation error. Replicated laboratory and field time series that jointly measure detectable DNA, total substrate-associated DNA and environmental DNA supply will be necessary to estimate these processes, quantify uncertainty and support quantitative ecological inference from passive eDNA samplers.

### 4.6 General conclusions

By distinguishing detectable adsorbed DNA from degraded, non-detectable DNA that may continue to occupy substrate capacity, this study shows that passive eDNA recovery can range from monotonic saturation to unimodal accumulation with post-peak decline. The analytical solutions demonstrate that detectable yield and the yield-maximising retrieval time are determined jointly by DNA input rate, degradation rate, residual retention of degraded DNA and substrate capacity.

Passive sampler yield is therefore not necessarily proportional to environmental DNA supply. Finite substrate capacity can compress quantitative differences among input conditions even under complete replacement, whereas non-replacement dynamics can further make recovery strongly dependent on retrieval time. Under prolonged deployment, a higher-input condition may yield less detectable eDNA than a lower-input condition, reversing the expected input-rate ranking. Similar adsorption and degradation dynamics among DNA sources may help preserve relative representation within a sampler, but this assumption requires empirical calibration.

In natural environments, competition from non-target DNA and material-specific substrate capacity may further influence the recovery of low-abundance targets, with implications for rare-species detection and metabarcoding. Deployment duration should therefore be treated as a central design parameter in passive eDNA sampling rather than a logistical detail. Time-series calibration across DNA input, total DNA load, degradation conditions and sampler materials will be necessary to identify suitable retrieval windows and support reliable quantitative interpretation.

## Acknowledgements

We are grateful to Dr. Takashi Kanbe and Mr. Kohei Nishitani for their comments and suggestions on the first draft. This study was supported by JSPS KAKENHI grant no. 23H00329, JSPS Program for Forming Japan’s Peak Research Universities (J-PEAKS) Grant Number JPJS00420230001, and the Environment Research and Technology Development Fund of the ERCA (JPMEERF20254M05) funded by the Ministry of the Environment of Japan.

## Author Contributions

Both authors contributed to the study conception and design. The first draft of the manuscript was written by HA, and both authors revised the manuscript.

## Declaration of generative AI in the manuscript preparation process

During the preparation of this work, the authors used GPT-5.6 (OpenAI) to conduct minor text editing. After using this tool, the authors reviewed and edited the content as needed and take full responsibility for the content of the published article.

## Data Archiving Statement

This study does not contain any data requiring archiving.

## Conflict of Interest

The authors have no competing interests to declare.

## Notes

### Competing Interest Statement

The authors have declared no competing interest.

